# Long noncoding RNA AL109754.1 Regulates Myeloid Dendritic Cell Differentiation and Potentiates TLR signaling

**DOI:** 10.1101/2023.03.01.530613

**Authors:** Raza Ali Naqvi, Araceli Valverde, Afsar Naqvi

## Abstract

Dendritic cells (DCs) are key antigen presentation cells (APC) that bridge innate and adaptive immune functions to contain the pathogenic threats. Long noncoding RNAs (lncRNAs) are implicated in functional regulation of various biological processes including inflammation and immunity. However, the knowledge on myeloid DC expressed lncRNA repertoire and their regulatory functions is limited. In this study, we have reconnoitered the time-kinetics of lncRNA expression profiles during monocyte-to-DC differentiation and their roles in shaping DC functions. Our RNA-seq data identified thousands of differentially expressed lncRNAs associated with primary human monocyte-to-DC differentiation *in vitro*. We selected two lncRNAs *viz*., AL109754.1 and AC093278.2 that were enriched during DC differentiation. Knockdown of AL109754.1 but not AC093278 affects DCs differentiation as observed by marked reduction of surface markers CD1a, CD93 and CD209. These DCs also exhibit significant reduction in the expression of TLR 2, 4, 5, 7 and 9, suggesting that AL109754.1 expression is critical in maintaining TLR expression in DCs. Furthermore, reduced phosphorylation of NF-κB, IRF3 and IRF7 in AL109754.1 knockdown DCs treated with TLR agonists further substantiate their role in potentiating TLR signaling. Mechanistically, AL109754.1 knockdown DC showed significant downregulation of multiple NF-κB-induced genes and time-dependent inhibition of pro-inflammatory cytokine (IL-1β, IL-6, IL-8 and TNFα) secretion upon challenge with TLR 4, 5, or 7 agonists. Overall, this study characterized novel functions of AL109754.1 that regulates DC differentiation, and TLR-dependent innate immune activation.

## Introduction

Dendritic cells (DCs) are professional antigen-presenting cells (APCs) and are the result of efficient and well-orchestrated lympho-myeloid hematopoiesis of bone-marrow-derived cells (1). Functionally, DCs recognize and capture the antigens derived from invading pathogens, process them into peptides, migrate towards lymph nodes and present them to T cells and finally, activate the specific adaptive immunity to clear the pathogenic threat (2–4). Therefore, DC differentiation and function in response to antigens should be tightly regulated to maintain immune homeostasis.

Recently, long noncoding RNAs (lncRNAs) have emerged as crucial regulator of transcriptome and has potential to modulate the biological behavior of the cells including immune functions (5, 6). LncRNAs are ≥ 200 nucleotides in length and do not possess intrinsic potential to encode for proteins like mRNA. Multiple genome-wide transcriptomic studies unravel the fact that ∼10-20 fold more genomic sequences encode for lncRNA compared to protein-coding genes (7). In this pursuit, we have recently deciphered the regulatory role of lncRNAs in innate immune functions and identified lncRNAs RN7SK and GAS5 in regulating macrophage polarization and functions (8). However, the role of lncRNAs in monocyte to DC differentiation is less explored, as only a handful of studies have suggested their putative roles in DC differentiation and function. Using high-throughput methods *viz*., transcriptome microarray analysis and RNA sequencing (RNA-seq), Wang et al. deciphered the role of STAT3-binding lncRNA in monocyte to DC differentiation (9). Lentivirus-mediated knockdown of this lncRNA impaired differentiation of monocyte to DCs, antigen uptake and presentation; and reduced proliferation of allogenic CD4+ T cells (10). Furthermore, Zhou et al., reported the differential expression of 962 lncRNAs in mature vs immature DCs in acute retinitis (AR) (10). *In vitro* treatment of human PBMCs with granulocyte-macrophage colony stimulating factor (GM-CSF) and interleukin-4 (IL-4) results in the downregulation of lncRNA HOTAIRM1 expression, which showed an inverse relationship with monocyte to DC differentiation (11). Though all these studies suggested the roles in lncRNAs in DC differentiation and function, in-depth time-dependent kinetics of lncRNAs expression is not well-studied so far. Therefore, exploring the lncRNA expressional dynamics at different time points of monocyte-to-DCs differentiation cells will provide the treasure trove of information that would be pivotal in understanding immune dysregulations pertaining to multiple diseases under distinct immune microenvironments.

DC functions toward pathogen clearance are facilitated by well-equipped multitude of pattern-recognition receptors (PRRs) involved in pathogen recognition (12). Toll-like receptors (TLR) 1-9 are the most widely studied and highly conservative family of PRRs across the species. Every TLR recognizes unique molecular patterns on pathogens (12) and the engagement of TLR with specific pathogen patterns activates a set of effector transcription factors *viz*., Nuclear factor kappa-light-chain-enhancer of activated B cells (NF-κB), Interferon Regulatory Factors (IRFs), or Mitogen-Activated Protein Kinases (MAPK) to induce the expression of pro-inflammatory cytokines, chemokines, and type I IFNs to neutralize the infectious threat (13).

LncRNA-mediated regulation of TLR expression and activity in DCs is an understudied topic and the reports are sparsely available either in the context of THP1 cells or macrophages. In a study, Carpernter et al. showed that TLR2 ligation results in the induction of 62 different lncRNAs. They demonstrated that differential expression of *lincRNA-Cox2*, *lncRNA-Ehd1* and *lncRNA-Lyn* following TLR2 and TLR4 activation (15). LncRNA *THRIL* regulates TNFα expression via interacting with hnRNPL during activation of THP1 macrophages (16). To date, only limited studies have examined TLR expression regulation in immune cells as reflection of lncRNA expressional dynamics.

In this systematic and comprehensive study, we have profiled time-kinetics of lncRNA expression during monocyte-to-DC differentiation using RNA-Seq. We have characterized the role of an unannotated lncRNA AL109754.1 in DC differentiation and its regulation of TLRs and their downstream signaling.

## Materials and Methods

### Primary human monocytes (PBMCs) isolation and differentiation

Freshly collected peripheral blood from healthy adults was used to isolate PBMCs by density gradient centrifugation using Ficoll Paque (GE Healthcare, Piscataway, NJ, USA) as previously described (17). CD14^+^ cells were isolated (>95% pure) using magnetic beads according to the manufacturer’s instructions (Miltenyi Biotec). Monocytes were treated with GM-CSF (50 ng/mL; both 50 ng/ml; PeproTech) and IL-4 (50 ng/mL; both 50 ng/ml; PeproTech) in RPMI supplemented with penicillin (100 U/ml), streptomycin (100 mg/ml) for 7 days to obtain DCs.

### RNA extraction, library construction and sequencing

Total RNA was extracted using miRNeasy kit (Qiagen, CA, USA) as per manufacturer’s instructions. Bioanalyzer 2100 and RNA 6000 Nano LabChip Kit (Agilent, CA, USA) were used to assess the RNA quality and quantity. RNA samples with RIN number >7.0 were processed for the RNA-Seq experiments. Firstly, we have removed ribosomal RNA according to Epicentre Ribo-Zero Gold Kit (Illumina, San Diego, USA). Then, using divalent cations under elevated temperature, the ribo-negative RNA fractions were fragmented into small pieces and reverse-transcribed for the creation of final cDNA library using strand-specific library preparation by dUTP method. 300±50 bp was the average insert size for the paired-end libraries. Afterwards, pair-end 2 X 150bp sequencing was done on illumina Hiseq 4000 platform following the vendor’s recommended protocol.

The removal of reads possessing adaptor contamination, low quality bases and undetermined base was done by Cutadapt (18) method and the sequence quality was checked by FastQC (http://www.bioinformatics.babraham.ac.uk/projects/fastqc/). Subsequently, the reads were mapped on human genome (version) by Bowtie2 (19) and Tophat2 (20), and then were assembled using StringTie (21) for every sample. All transcriptomes from various samples were merged and a comprehensive transcriptome was reconstructed using perl scripts and gffcompare (https://github.com/gpertea/gffcompare/). Once the transcriptome was generated, we have estimated the expression levels of all transcripts using StringTie and Ballgown (21). After that, lncRNA identification process was performed, wherein, the transcripts mapped to mRNAs, and transcripts shorter than 200 bp were discarded. Then, the transcripts showing CPC score <-1 and CNCI score <0 were removed and the remaining transcripts aligned with the genome were considered as lncRNAs (22). FPKM (FPKM = [total_exon_fragments / mapped_reads (millions) × exon_length (kB)]) were used to evaluate the expression level for lncRNAs using StringTie (21).

### Quantitative RT-PCR analysis

Monocytes and monocyte-derived DC were gently washed with PBS in the culture plate. After that, 700 µl TriZol reagent (Invitrogen, CA, USA) was added to each well and RNA was isolated using miRNeasy micro kit (Qiagen, Gaithersburg, MD, US). 250 ng RNA was used to synthesize cDNA, by high capacity cDNA reverse transcription kit (ThermoFisher Scientific, Grand Island, NY, USA). The expression levels of lncRNAs AL109754.1 and AC093278.2 were analyzed by qPCR reaction using SYBR Green Gene Expression Master Mix (Applied Biosystems, USA) in a StepOne 7500 thermocycler (Applied Biosystems, USA). To calculate ΔCt, the cycle number of experimental conditions and their controls were first normalized by their corresponding Ct values of β-actin gene. The Ct values of three replicates were analyzed to calculate fold change using the 2^−ΔΔCt^ method. Sequences of designed primers are provided in **Supplemental Table I**.

### Transient siRNA Transfection

DCs were transfected with control or lncRNA-specific siRNA using Lipofectamine 2000 (Invitrogen-Life Technologies Corporation, Carlsbad, CA, USA), as per the manufacturer’s instructions. Two siRNAs specific to AL109754.1 and AC093278.2 were designed using online tool (IDT) (Supplemental Table II). Control siRNA were also purchased from IDT. (Gaithersburg, MD, USA).

### Flow Cytometry

Cells were washed two times with PBS supplemented with 1% (v/v) BSA, treated with enzyme free cell dissociation buffer (Gibco). Harvested cells were washed in PBS-1% BSA two times to remove the traces of cell dissociation buffer. For DC differentiation experiment, the cells were stained with anti-CD1a (FITC anti-human CD1a Antibody, clone: HI149, BioLegend), anti-CD93 (PE conjugated anti-human CD93 Antibody, clone: VIMD2, BioLegend), anti-CD209 (APC conjugated anti-human CD209 Antibody, clone: 9E9A8, BioLegend). For TLR staining experiments, cells were stained with TLR1 (BV421 conjugated Mouse Anti-Human CD281, clone: GD2.F4, BD Biosciences), TLR2 (FITC conjugated anti-human CD282 Antibody, clone: W15145C, BioLegend), TLR3 (Brilliant Violet 711™ conjugated anti-human CD283, clone: TLR-104, BioLegend), TLR4 (PE conjugated anti-human CD284, clone: HTA125, BioLegend), TLR5 (Alexa Fluor® 488 conjugated anti-human CD285 Antibody, clone: S16021I, BioLegend), TLR6 (PE conjugated anti-human CD286 Antibody, clone: TLR6.127, BioLegend), TLR7 (Alexa Fluor® 700-conjugated Antibody, clone: 533707, R&D Systems), TLR8 (PE anti-human CD288 Antibody, clone: S16018A, BioLegend) and TLR9 (APC conjugated anti-human CD289 Antibody, clone: S16013D, BioLegend) antibodies. Afterwards, the cells were washed 2 times with PBS-1% BSA. All samples were analyzed in CytoFLEX S Flow Cytometer. FlowJo_v10.8.1 software (Tree Star, Ashland, OR) was used to analyze the flow cytometry data.

### TLR stimulation and cytokine production

To evaluate the TLR-mediated downstream phosphorylation, day 6 DCs were cultured overnight (18 hours) in RPMI-1640 with TLR agonists (Human Toll-like receptor (TLR) 1– 9 kit; InvivoGen). Following concentrations of TLR agonists were used to stimulate TLR2 (HKLM: 10^8^ cells/mL), TLR3 (1 µg/ml), TLR4 (LPS-EK standard, 10 ng-10 µg/ml], TLR5 (FSL-1: 10 ng-10 µg/ml), TLR7 (imiquimod: 10 ng-10 µg/ml), TLR8 (ssRNA40/ LyoVec™^;^ 5µg/ml) and TLR9 (ODN2006: 5µM) in monocyte-derived DCs. Culture supernatants were collected after 4h and 18h time intervals, and were analyzed for cytokine levels by Milliplex Luminex bead-linked immunoassay, according to the manufacturer’s protocol (EMD Millipore). All culture supernatants obtained from TLR stimulation of DCs culture and standards (provided in the kit) were run in duplicate during the assay and the measurements were normalized to baseline values of cytokines obtained from unstimulated cells.

### Phosphoflow

To obtain the phosphorylation states of downstream TLR signaling effector transcription factors (TFs), the DCs were fixed post-TLR stimulation at 4°C for 30 min. Harvested cells were washed two times with RPMI-1%BSA and fixed with Cytofix Buffer (BD Biosciences). After permeabilization, DCs were washed two times with RPMI-1%BSA and incubated with antibodies. Following antibodies were used to detect the downstream phosphorylation: NF-κB (BV421 conjugated NF-κB [p65], clone: K10-895.12.50, BD Biosciences), IRF7 (Alexa Fluor® 647 conjugated Mouse anti-IRF-7 [pS477/pS479], clone: K47-671, BD Biosciences), IRF3 (PE conjugated Rabbit mAb Phospho-IRF-3 [Ser396], clone: D6O1M, Cell Signaling).

### PCR array

Monocyte-derived DCs transfected with AL109754.1 siRNAs were challenged with TLR2, TLR5 or TLR9 agonists. After 24h post-challenge, the cells were washed gently with PBS and Qiazol was added to isolate total RNA. Gene expression was examined using PCR array plate containing 88 primer sets directed against human NF-κB pathway genes and 8 housekeeping primer sets (Real Time Primers, LLC Elkins Park, PA). Briefly, 1µg of total RNA was used for cDNA synthesis and real-time PCR was performed on StepOne 7500 thermocycler (Applied Biosystems, Carlsbad, CA). The data was analyzed using the Qiagen GeneGlobe Data Analysis Center (Source: https://www.qiagen.com/us/shop/genesand-pathways/data-analysis-center-overview-page/). B2M was used as housekeeping to normalize the expression levels.

### Statistical Analysis

Data were analyzed and plotted using GraphPad Prism (La Jolla, USA). The results are represented as SD or SEM from independent replicates (n=3-6/group). *P*-values were calculated using Students t-test (non-parametric), and the *value* <0.05 was considered significant (\**p < 0.05, **p < 0.01, ***p < 0.001)*.

## Results

### Time-kinetics of global lncRNA expression profiles during monocyte-to-DC differentiation

To characterize the in-depth lncRNA profiles during monocytes to DC conversion, we isolated RNA from primary CD14^+^ human monocytes (n=5; healthy donors) and from monocytes treated with GM-CSF & IL-4 at different time points: 18h (n=4), day 3 (n=4), day 5 (n=4) and day 7 (n=5). Total RNA was then subjected to poly-adenylated paired-end RNA sequencing to examine lncRNA expression kinetics during monocytes-to-DCs conversion *in vitro*. We obtained 466 million (monocytes), 457 million (18h), 259 million (day 3), 508 million (day 5) and 547 million (day 7) raw reads (**Table I**). After quality control, ∼380, 377, 311, 420, 505 million of clean reads were obtained for monocytes, and the differentiating DCs (**Table I**). Approximately 75-79% reads uniquely mapped to the human genome (GRCh38.79) in all the samples. All the transcripts exhibiting overlap with known mRNAs and the transcript shorter than 200 bp were discarded. Other properties, such as transcript lengths, length of open reading frame (ORF), and exon number of lncRNAs in every condition were also analyzed (**Supplemental Fig. S1A-C**) Afterwards CPC (Coding Potential Calculator) (*25*) and CNCI (Coding-Non-Coding Index) (*26*) were used to envisage the coding potential of the transcripts. All transcripts with CPC score <-1 and CNCI score <0 were also removed and all the remaining transcripts with class code j (at least one splice junction is shared with a reference transcript), i (a transfrag falling entirely within a reference intron), o (generic exonic overlap with a reference transcript) and x (exonic overlap with reference on the opposite strand) were considered as known lncRNAs. Approximately, each class of lncRNAs group were represented by 3% (j class), 70% (I class), 4% (o class), and 3% (x class) lncRNA transcripts (**Supplemental Fig. S2)**. In addition, we have also identified novel lncRNA sequences in our dataset categorized as unknown (class u) (*Naqvi et al., unpublished data*).

**Table I:** Summary of RNA-seq data showing total, filtered and mapped reads in various samples.

| Sample | Raw data | Valid reads | Mapped reads | Unique Mapped reads |
| --- | --- | --- | --- | --- |
| Mon_1 | 102891216 | 86600654 | 52196486(60.27%) | 42388685(48.95%) |
| Mon_2 | 92418804 | 72499094 | 54132184(74.67%) | 44763582(61.74%) |
| Mon_3 | 104487562 | 85099740 | 55840893(65.62%) | 45310437(53.24%) |
| Mon_4 | 93634330 | 78266364 | 56516789(72.21%) | 45188912(57.74%) |
| Mon_5 | 72587344 | 58440778 | 38680726(66.19%) | 29822980(51.03%) |
| DC1_18h | 128124302 | 101575394 | 71642246(70.53%) | 55169278(54.31%) |
| DC2_18h | 110397350 | 90925258 | 64786245(71.25%) | 52778473(58.05%) |
| DC3_18h | 94993928 | 79676582 | 55450841(69.59%) | 45437577(57.03%) |
| DC4_18h | 124134500 | 105067770 | 81003162(77.10%) | 66642107(63.43%) |
| DC1_D3 | 121085682 | 97928812 | 75894192(77.50%) | 61992558(63.30%) |
| DC2_D3 | 86331148 | 70867188 | 62788086(88.60%) | 50865793(71.78%) |
| DC3_D3 | 84553250 | 70786566 | 65078564(91.94%) | 54106585(76.44%) |
| DC4_D3 | 88408552 | 71834318 | 62679356(87.26%) | 51417765(71.58%) |
| DC1_D5 | 120867264 | 99806534 | 71260577(71.40%) | 56893915(57.00%) |
| DC2_D5 | 137195650 | 110884424 | 79349557(71.56%) | 62301859(56.19%) |
| DC3_D5 | 119677550 | 96849298 | 52585983(54.30%) | 41128107(42.47%) |
| DC4_D5 | 130538136 | 104888006 | 60550008(57.73%) | 48051523(45.81%) |
| DC1_D7 | 134414176 | 98819894 | 71837523(72.70%) | 56445967(57.12%) |
| DC2_D7 | 124507688 | 101439508 | 62424861(61.54%) | 49453102(48.75%) |
| DC3_D7 | 144368158 | 107142740 | 90611473(84.57%) | 71665770(66.89%) |
| DC4_D7 | 139449488 | 87806720 | 73491762(83.70%) | 57556812(65.55%) |
| DC5_D7 | 139464464 | 110612090 | 67780862(61.28%) | 53060422(47.97%) |

The time kinetics of lncRNA profiles during DC differentiation process were analyzed with a cutoff range between -1.25 and 1.25 with *p* value <0.05. Heatmaps shown in **Figure 1 (A-D)** clearly demonstrate that lncRNA expression changes precedes DC lineage commitment. Compared with monocytes, we identified a large number of differentially expressed lncRNAs at 18h, 3d, 5d and 7d post-differentiation suggesting that a unique repertoire of lncRNA are expressed during monocyte-to-DC differentiation (**Fig. 1A-D**). More precisely, following numbers of known lncRNAs were found to be differentially expressed (up- and down-regulated) during monocytes to DC conversion at 18h, 3d, 5d and 7d: 1251 (738 up and 513 down), 1344 (705 up and 639 down), 1767 (1029 up and 738 down) and 793 (326 up and 467 down), respectively (**Fig. 1E**). We noticed that ∼17% of lncRNAs were common across all the time-points, while ∼12, 10, 9 and 16% lncRNAs were unique to 18h, day 3 day 5 and day 7, respectively.

**FIGURE 1:**
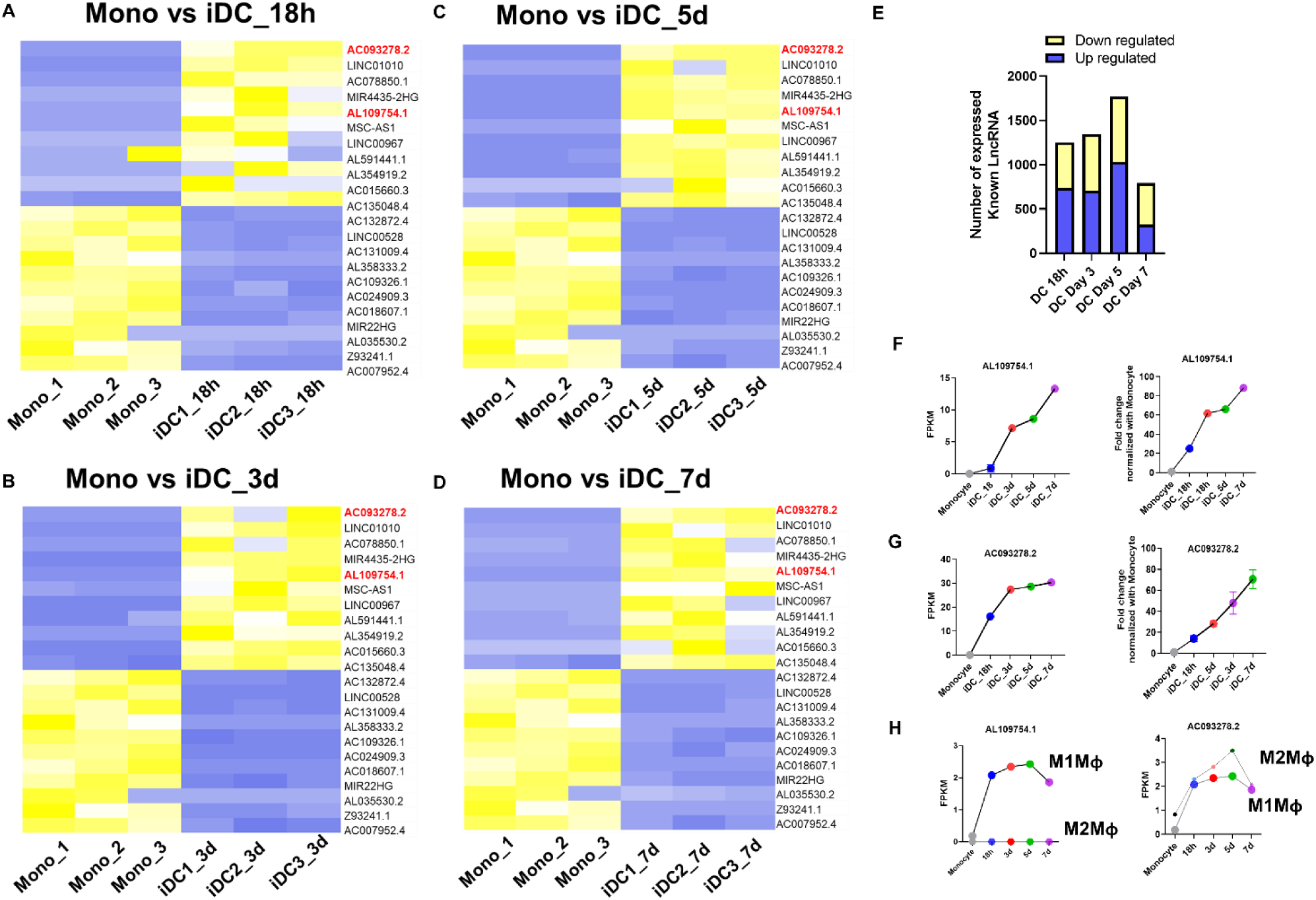
LncRNA expressional dynamics during monocyte-to-DC differentiation by RNA-sequencing. (A-D) Time kinetics of known lncRNAs in monocyte-to-DCs differentiation in 18h, 3d, 5d and 7d obtained from RNA-seq data. The heap map was generated after removal of ≤ transcripts shorter and confirmation of their coding potential by CPC (Coding Potential Calculator) and CNCI (Coding-Non-Coding Index). LncRNA was identified on following criterion: CPC score <-1 and CNCI score <0. **(E)** Evaluation of number of up and downregulated lncRNAs based on fold change (a cut off between - 1.25 and 1.25) and p ˂ 0.05 (multiple ANOVA) each condition. (**F**) Expressional validation of time dependent kinetics of lncRNA AL109754.1 during monocyte to DC differentiation using RT-qPCR analysis. (**G**) Expressional validation of time dependent kinetics of lncRNA AC093278.2 during monocyte to DC differentiation using RT-qPCR analysis. Time dependent expressional kinetics of during monocyte to DCs differentiation. (**H**) Evaluation of AL109754.1 and AC093278.2 expression in M1 and M2 macrophages. In RT-qPCR analysis each data point is representative of at least three different experiments. Non parametric *t-tes*t was used to evaluate statistical significance using GraphPad 9. *p < 0.05, **p < 0.01, ***p < 0.001.

To validate the RNA-seq data, we selected two DC-enriched lncRNAs viz., AL109754.1 and AC093278.2 and examined their expression by RT-qPCR in an independent cohort (**Fig. 1 F**). Our candidate lncRNAs showed time-dependent induction in their expression during DC differentiation and corroborate with the RNA-seq data. Together, our results demonstrate that thousands of lncRNAs are involved in monocyte-to-DC differentiation suggesting their functional regulatory role in this biological process. We next asked whether these lncRNAs are expressed in other monocyte-derived cells viz., M1 and M2 polarized macrophages. Interestingly, AL109754.1 expression in M1Mφ exhibit similar expression pattern of upregulation as monocytes-derived DCs but was not detected in M2Mφ, thereby suggesting the pro-inflammatory nature of AL109754.1 (**Figure 1G**). On the contrary, AC093278.2 was detected and showed time-dependent increase in both M1 and M2 macrophages (**Figure 1G**). However, the FPKM levels of both AL109754.1 and AC093278.2 in DC were ∼10-fold higher than either polarized Mφ indicating a key role of these lncRNAs in myeloid DC differentiation and/or innate immune functions.

### LncRNA AL109754.1 regulates the differentiation of myeloid dendritic cells

LncRNAs regulate differentiation of various immune cells by integrating extracellular signals with multitude of biological pathways, which eventually facilitate immune cells to adapt and modulate their functions aptly in response to rapid changes in the microenvironment (23). Our RNA-seq data showed upregulation of AL109754.1 and AC093278.2 during monocyte- to-DC differentiation. We next asked whether knockdown of DC-enriched AL109754.1 and AC093278.2 regulate the differentiation process.

We designed two siRNAs for each candidate lncRNA to examine their potency to knockdown target RNA. DC were transfected with individual siRNAs or their combination to assess RNAi efficacy. Our results show that siRNA A and siRNA B transfection significantly reduced the levels of their target lncRNA AL109754.1 and AC093278.2, respectively (**Supplemental Fig. 3, Supplemental Table II)**. Day 3 DCs were transfected with lncRNA targeting siRNAs and their impact on DC differentiation was examined by surface markers using multi-parametric flow cytometry. Previous study from our laboratory showed the induction of CD1a, CDw93, and DC-SIGN (CD209) during monocyte-to-DC differentiation (24). Compared to scramble, histograms and scatter plots show that lncRNA AL109754.1 knockdown remarkably reduced the expression of CD1a (44.5±3% vs 25.6±12.5%), CD209 (90.0±2.5% vs 67.3±4.6%) and CDw93 (87.3±6.6% vs 71.4±13.5%) (**Fig. 2A, 2B, 2C; left and middle panels**). Knockdown of AC093278.2 did not cause any significant change in the levels of these markers. We also quantified the expression of these differentiation markers by calculating geometric mean florescence intensity (Geo. MFI). CD1a (78.5±6.55%; *p<0.005*), CD209 (75.81±12.2%; *p<0.005*) and CDw93 (64.1%±13.7; *p<0.005*) levels show significant downregulation in AL109754.1 knockdown DCs, while AC093278.2 knockdown did not affect the expression of tested surface markers (CD1a: 92.9±5.16.55%; CD209: 119.2±10.32%; CDw93: 85.13%±15.24) (**Fig. 2A-C, right panel**). Together, our results clearly suggest that the enrichment of lncRNA AL109754.1 is critical in monocyte-to-DC differentiation process.

**FIGURE 2:**
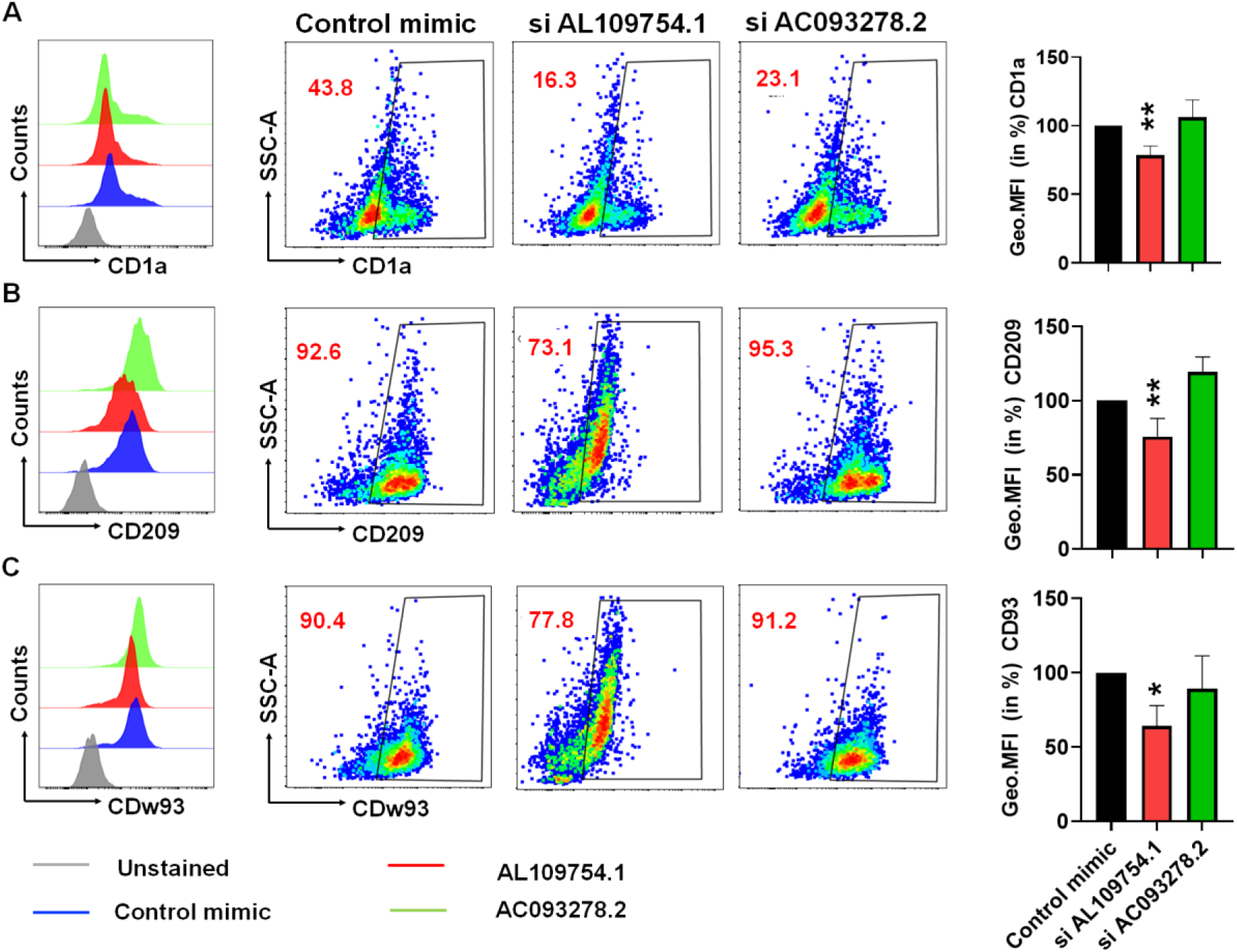
LncRNA AL109754.1 regulates the differentiation of monocyte-to-DC. CD14+ purified monocytes derived from healthy human donors (n=9/group) were induced to DC via GM-CSF and IL-4 treatment. Day 3 differentiated DC were transfected with siRNAs targeting lncRNAs AL109754.1 and AC093278.2 and the expression of DC markers **(A)** CD1a**, (B)** CD208, and **(C)** CDw93 were assessed by flow cytometry analysis. Half overlaid histograms were generated in Flowjo_v10.8.1 to elucidate the expressional comparison of DC makers in control and candidate lncRNA knockdown. Different colors in half overlaid histograms showing experimental conditions. Middle panel in (**A-C**) show representative scatter plots depicting percentages of DC differentiation markers. Right panel shows normalized geo. MFI percentages. Each bar shows mean ±SD value. Student’s t-test were used to calculate p-values and p <0.05 was considered significant. *p < 0.05, **p < 0.01, ***p < 0.001.

**FIGURE 3:**
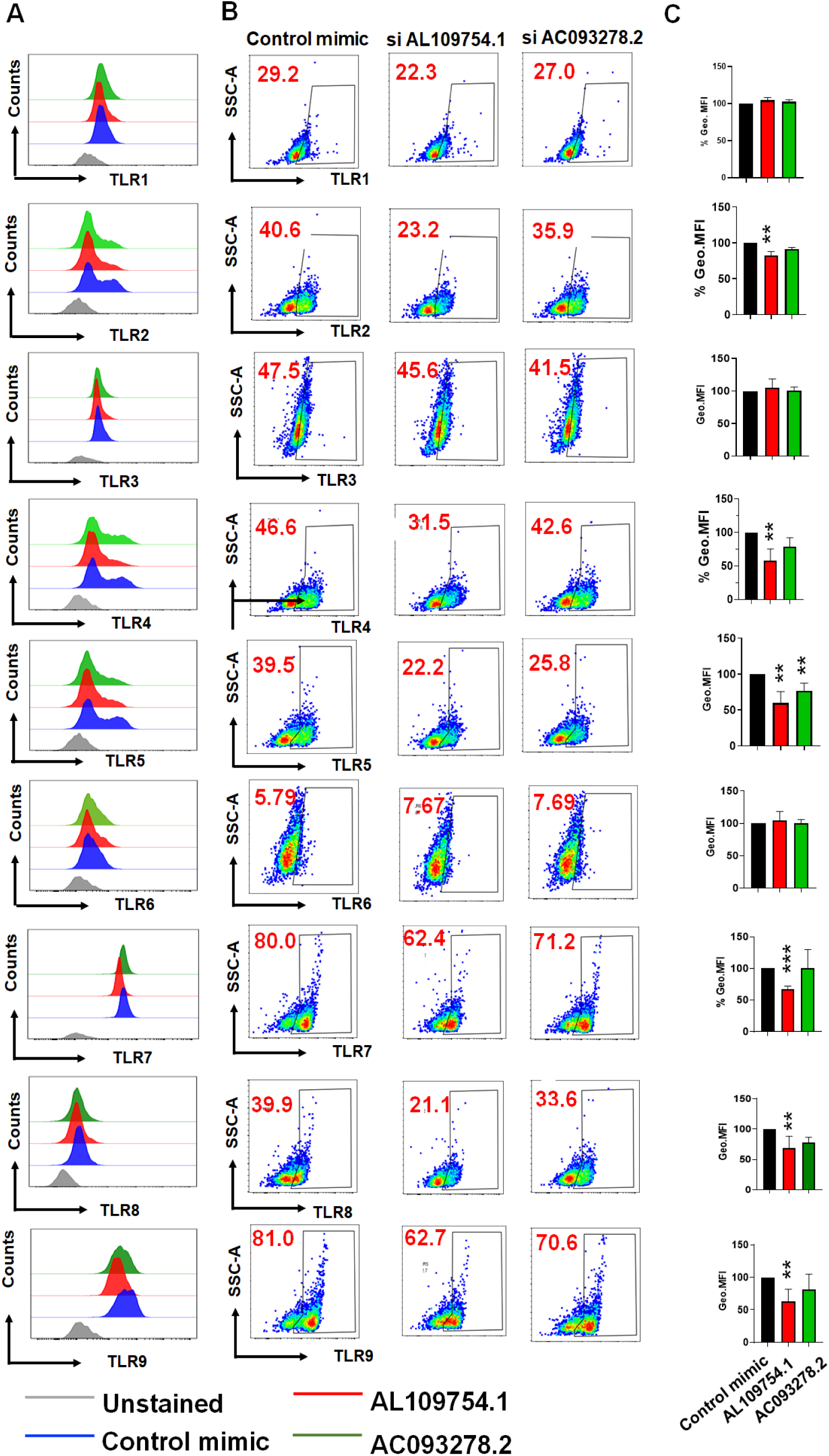
LncRNA AL109754.1 regulates TLR expression during the process of monocyte-to-DC differentiation. Monocyte-derived DC were transfected with siRNA (50nM concentration) targeting lncRNAs AL109754.1, AC093278.2 or control using lipofectamine 2000. The expression of TLRs (1–9) was evaluated 48 h post-transfection by flow cytometry. (**A)** Half overlaid histograms shows comparative expression of TLRs (1–9) in scramble and lncRNA knockdown DCs. Various colors indicate different experimental conditions. **(B)** Scatter plots showing the percentage of TLR expression on DCs upon lncRNA knockdown. **(C)** Normalized percent geometric MFI (Geo. MFI) of each TLR experiment. Geo. MFI values were calculated by normalizing with scramble and presented as percentages. Half laid histograms, and scatter plots were generated using FlowJO_v10.8.1. Same software was also used to calculate geo. MFI. The data present here was obtained from n=9 healthy individuals. Each bar shows mean ±SD. Students t- test were used to calculate p-values and P <0.05 was considered significant. *p<0.05, **p<0.01, ***p<0.001.

### Knockdown of lncRNA AL109754.1 reduces the TLR expression on DCs

TLRs are considered as one of the most prominent PRRs that recognize a wide-spectrum of antigens by the DCs (25–27). We next asked whether differentially-induced lncRNAs AL109754.1 and AC093278.2 regulate TLR expression pattern in DCs. We transfected day 3 DCs with siAL109754.1, siAC093278.2 or scramble and the expression of various TLRs was examined by flow cytometric analysis.

Our results showed differential impact of AL109754.1 and AC093278.2 knockdown on various TLRs; however, AL109754.1 RNAi robustly altered multiple extracellular and endosomal TLR expression. Summary of the TLR expression changes in lncRNA knockdown is presented in ***Table II***. We did not observe evident changes in TLR1 expression in either of the lncRNA knockdown (**Fig. 3A, left, middle panel**). The observed normalized MFI values were 104.5±3.7% and 102.6±2.5% for AL109754.1 and AC093278.2 knockdown, respectively (**Fig. 3A right panel**). The knockdown of AL109754.1 (82.7±4.2%; p<0.005) but not AC093278.2 (91.3±2.0%) significantly reduced TLR2 expression compared to scramble (**Fig. 3B, left and middle panel**). The expression of TLR4 was significantly affected by the knockdown of AL109754.1 (57.8±17.3%; p<0.005), but no changes were observed for AC093278.2 (78.5%± 13.9) RNAi (**Fig. 3C, left and middle panel**). Interestingly, TLR5 expression was modulated by the knockdown of both AL109754.1 and AC093278.2 (**Fig. 3D, left and middle panel**). Compared to scramble, TLR5 expression was downregulated in AL109754.1 and AC093278.2 knockdown DCs were 59.5±16.1% (p<0.005) and 76.9±10.5% (p<0.005), respectively (**Fig. 3D, right panel**). TLR6 expression was unaffected by lncRNA AL109754.1 (104.8±13.5%) or AC093278.2 (100.3±5.7%) knockdown (**Fig. 3E, left, right** and middle panel).

**Table II:**
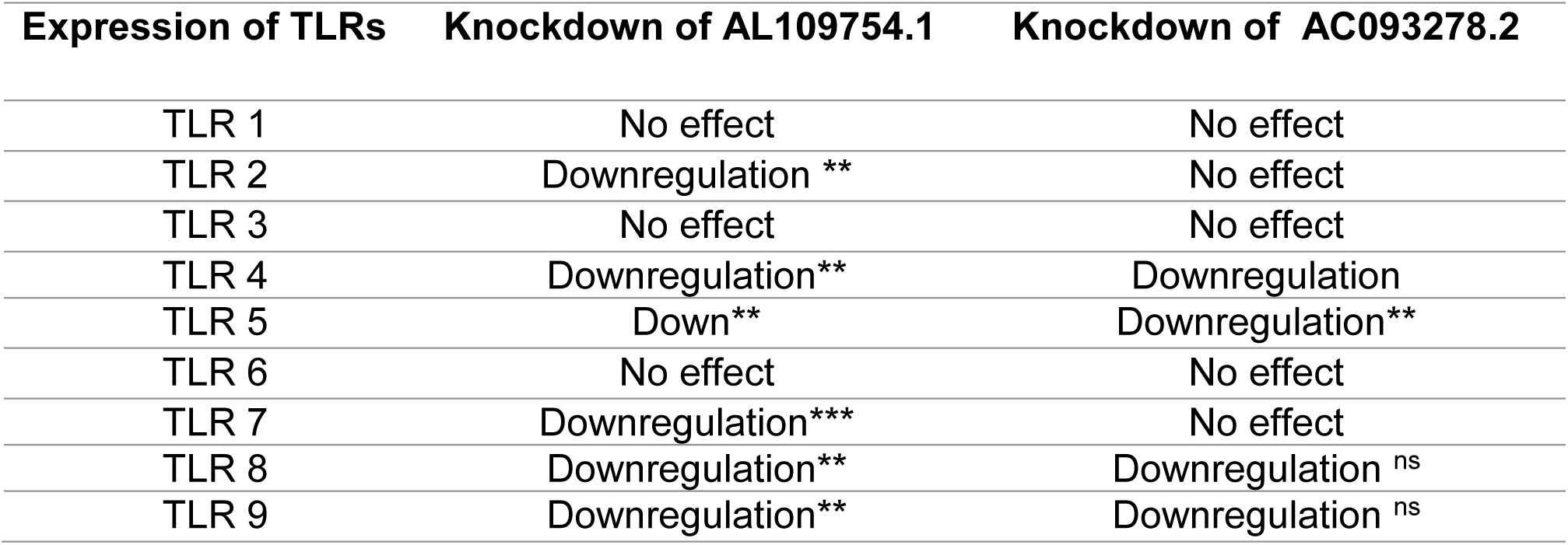
Effect of lncRNA knockdown on the expression of TLRs in DC. *p<0.05; **p<0.005;*** p<0.0005, ns: non-significant

Amongst the intracellular TLRs, TLR7, 8 and 9 but not TLR3 expression was impacted by lncRNA knockdown. The levels of TLR3^+^ cells in lncRNAs AL109754.1 (99.5± 5.9%) or AC093278.2 (99.8±5.1%) knockdown DC were comparable to scramble (**Fig. 3F, left, right and middle panel**). The expression of TLR7 was reduced by the knockdown of AL109754.1 (67.1±5.0%; *p<0.0005*), but siAC093278.2 had no impact (100.9±29%; **Fig. 3G, left and middle panel**). TLR8 levels showed reduction in AL109754.1 (68.7±19.2%; *p<0.005*) and AC093278.2 (77.3±8.9%) knockdown (**Fig. 3H, left and middle panel**). DC transfected with si-AL109754.1 showed reduced TLR9 levels (63.28±18.3%; *p* <0.005), while no significant changes were observed for siAC093278.2 (81.2±23.3%) (**Fig. 3I, left and middle panel**). Overall, these results clearly suggest that expression of lncRNA AL109754.1 in DCs regulate the expression of multiple plasma membrane (TLR2, TLR4, and TLR5) and endosomal (TLR7, TLR8 and TLR9) TLRs suggesting its essential role in DC maturation and innate immune responses.

### Pro-inflammatory cytokine secretion is regulated by lncRNA AL109754.1 in TLR agonist challenged DCs

TLR stimulation activates downstream signal transduction pathways in DCs to induce multiple proinflammatory cytokines, chemokines, and type I IFNs genes that eventually protect the host from the pathogenic insult (25–27). Next, we addressed whether TLR expression reduction upon lncRNA AL109754.1 knockdown alter the cytokine secretion capability of myeloid DCs, an essential component to provide adequate microenvironment for the DC maturation (28, 29). DCs transfected with siAL109754.1 were stimulated with ligands for TLR 2, 4, 5, 7 and 9, which showed significant changes in AL109754.1 knockdown DCs, and the supernatant were collected at 4h and 18h post-treatment for cytokine measurement.

Stimulation of siAL109754.1 transfected DCs with various TLR agonists showed impaired cytokine profiles in their culture supernatant (**Fig. 4**). In TLR2 stimulated DCs, we did not notice any decrease in IL-1β and TNFα levels in siAL109754.1 transfected cells at both time points (4h and18h). The effect of AL109754.1 knockdown on the decrease of IL-6 and IL-8 secretion was more pronounced at 18h and 4h (p<0.05), respectively. Interestingly, upon TLR4 stimulation siAL109754.1 transfected DCs showed significant reduction of IL-1β and IL-6 at 18h only (**Fig. 4**). Furthermore, IL-8 and TNFα levels were observed to be reduced at early (4h; p<0.05) and later (18h; p<0.05) time points, respectively. TLR-5 stimulated si-AL109754.1 transfected DCs showed reduced levels of IL-1β, IL-6 and TNFα at both time points, however effect of IL-6 reduction was more obvious at 18h (p<005). No significant changes were observed for IL-8. Interestingly, stimulation of transfected DCs with TLR7 showed reduction in all tested pro-inflammatory cytokines at both time points suggesting a central role of AL109754.1 in regulating TLR7 signaling. Likewise, IL-1β and TNFα also both showed reduction in siAL109754.1 transfected DCs stimulated with TLR9 agonist both at 4h and 18h time points. In this case, the potent reductions in IL-6 was seen at 18h (p<0.005), whilst the reduction of IL-8 was more significant at 4h (p<0.0005). However, we have also tested the effect of anti-inflammatory cytokine; IL-10, as a negative feedback regulation of inflammation. IL-10 was found reduced in all cases except stimulation of transfected DCs with TLR5 agonists (**Fig. 4**). Table III summarizes changes in cytokine levels upon TLR stimulation of si-AL109754.1 transfected DCs compared to control. Together, our results demonstrate that lncRNA AL109754.1 regulates TLR-stimulated cytokine secretion in DCs.

**FIGURE 4:**
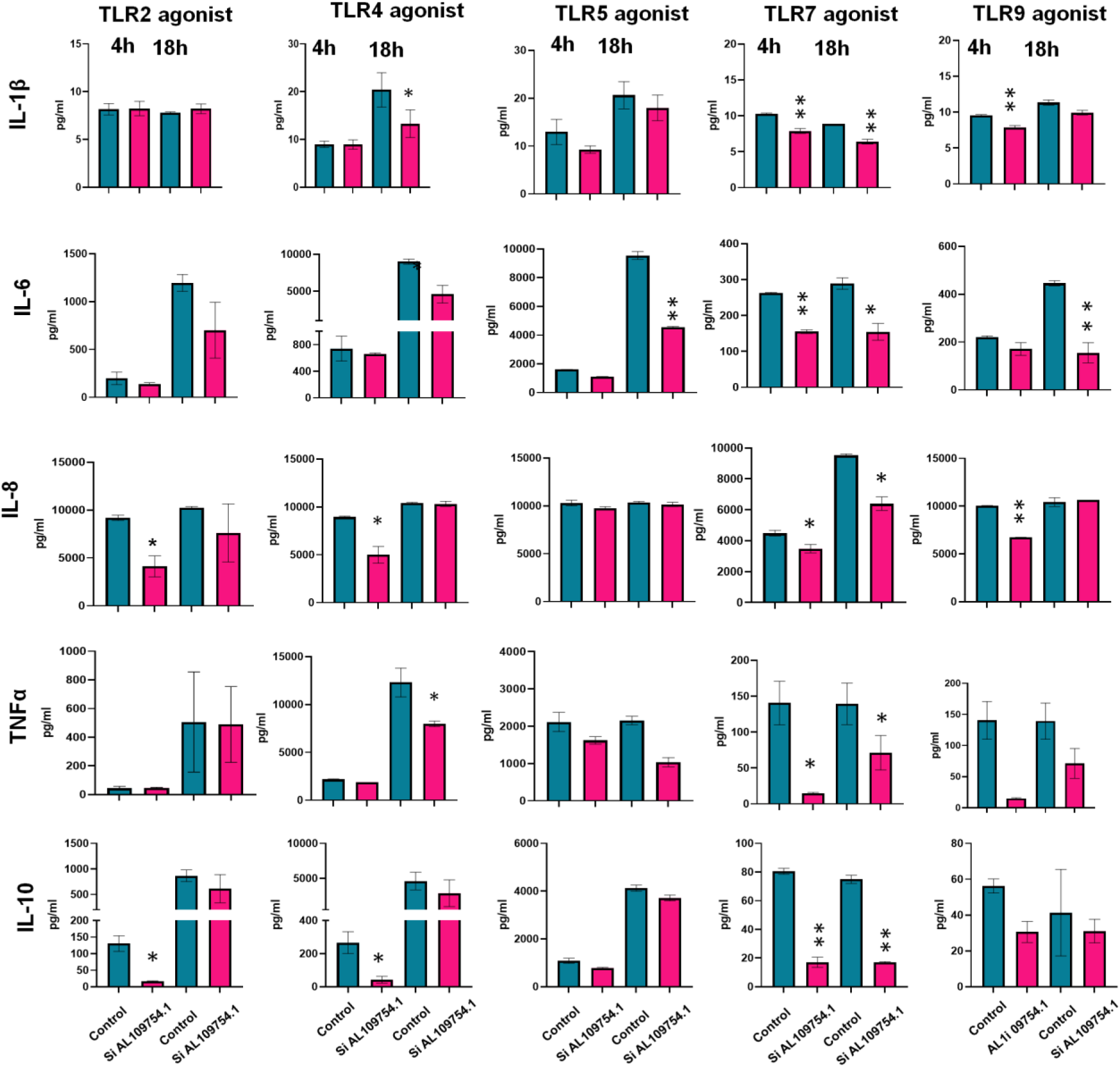
LncRNA AL109754.1 knockdown impairs pro-inflammatory cytokine secretion by DCs upon TLR stimulation. DCs were transfected with si-AL109754.1 and treated with various TLR agonists. The culture supernatants were collected after 4h and 18h time points and evaluated for cytokine analyses using Luminex MAGPIX® Multiplex system. All the bar graphs were generated in GraphPad prism 9. Each bar shows mean ±SD. Students t-test were used to calculate P-values and P<0.05 was considered significant. *p < 0.05, **p < 0.05, ***p < 0.005. All figures represent the experiments done with n=4 healthy donors.

**Table III:** Secretion of pro-inflammatory cytokines in AL109754.1 knockdown DCs upon TLR stimulation. NE: No effect, ↓: Reduced cytokine levels in culture supernatants; *p<0.05; **p<0.005;*** p< 0.0005

| | IL-1β | | IL-6 | | IL-8 | | TNFα | | IL-10 | |
| --- | --- | --- | --- | --- | --- | --- | --- | --- | --- | --- |
|  | 4h | 18h | 4h | 18h | 4h | 18h | 4h | 18h | 4h | 18h |
| TLR 2 | NE | NE | NE | ↓ | ↓* | ↓ | NE | NE | ↓* | ↓ |
| TLR 4 | NE | ↓ | ↓* | ↓* | ↓** | NE | ↓* | NE | ↓* | ↓ |
| TLR 5 | ↓* | ↓* | ↓*** | ↓** | NE | NE | ↓* | ↓* | ↓* | ↓* |
| TLR 8 | ↓*** | ↓** | ↓** | ↓* | ↓* | ↓** | ↓* | ↓* | ↓* | ↓* |
| TLR 9 | ↓* | ↓ | NE | ↓** | ↓*** | NE | ↓ | ↓*** | ↓* | ↓ |

### AL109754.1 regulates the downstream TLR-dependent phosphorylation

Pathogen recognition by TLRs triggers a signaling cascade, which involves recruitment of a variety of cytoplasmic proteins that eventually activate multiple transcription factors (NF-κB, IRF, etc.) via phosphorylation (30, 31). However, each TLR might phosphorylate different transcription factors/proteins to elicit unique immune response. Having confirmed that the knockdown of AL109754.1 affects multiple TLR expression during monocyte to DC differentiation and control innate immune cytokine secretion, we next evaluated the downstream phosphorylation cascade in monocytes-derived DCs.

Recent reports have elucidated the significance of IRF3 in the TLR4 signaling events (32). Furthermore, Fitzgerald et al. has shown that TLR4/MD2 expressing HEK increases the activation of exogenously expressed IRF7 upon stimulation with LPS (33). Therefore, we stimulated siAL109754.1 transfected DCs with TLR4 agonist *E. coli* LPS and evaluated the phosphorylation of IRF3, IRF7 and NF-κB (p65) 18h post-stimulation using phosphoflow. We observed significantly reduced pIRF3 (95.4±3.4% vs 47.5±4.4% vs; *p<0.005*) and pIRF7 (70.8±2.67% vs 46.1±5.5% vs %; *p<0.05*) positive cells in AL109754.1 knockdown DCs compared to control siRNA. However, the phosphorylation of p65^+^ (NF-κB) did not show significant change in AL109754.1 knockdown (56.3±3.1% vs 54±2.1%) DCs (**Fig. 5A, Left, middle and right panel**).

**FIGURE 5:**
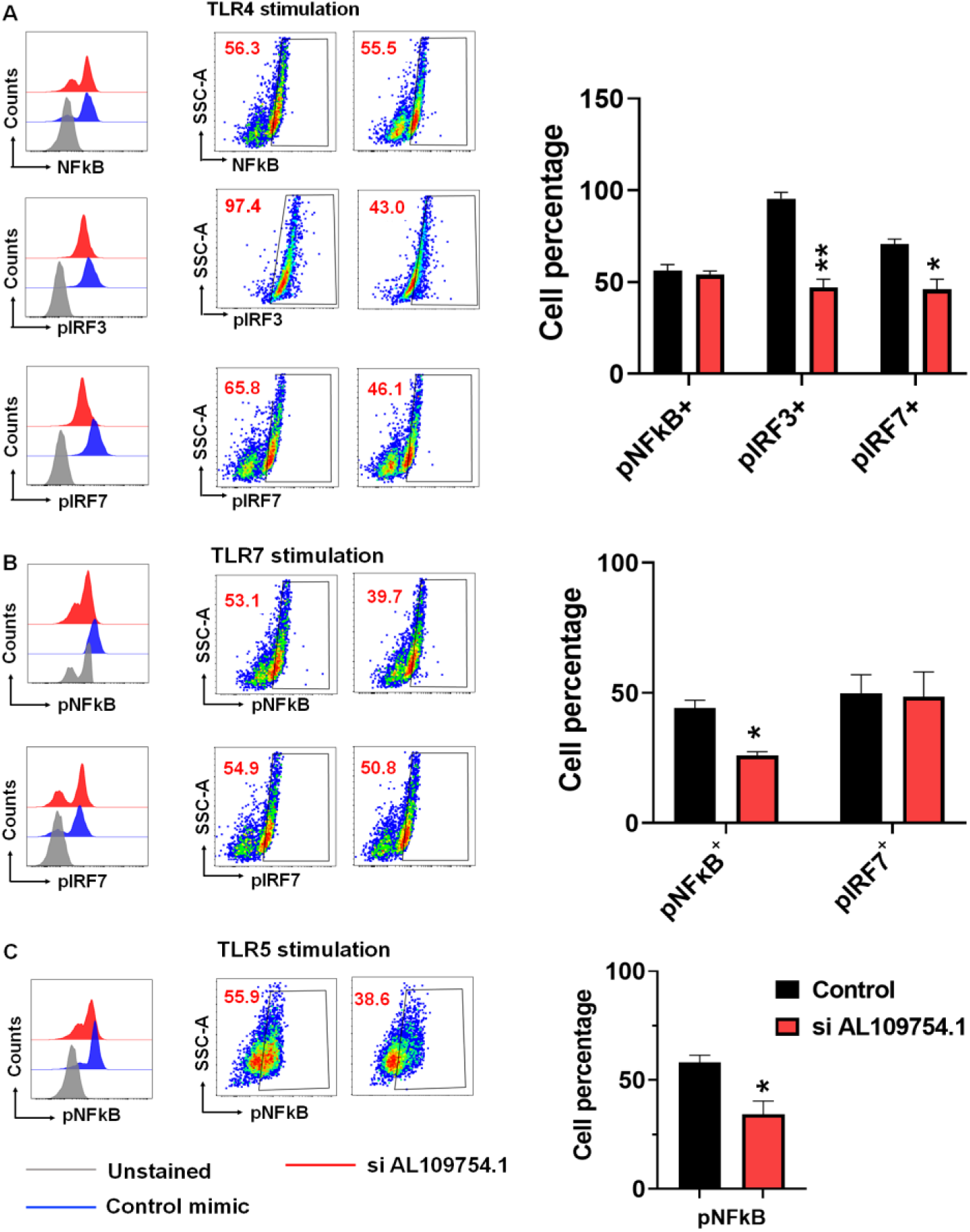
LncRNA AL109754.1 regulates TLR signaling dependent phosphorylation. *In vitro* human monocyte-derived DC were transfected with scramble or si-AL109754.1. After 48h, cells were challenged with TLR2, TLR4, TLR5, TLR7 or TLR9 agonists. (**A**) Effect of lncRNA AL109754.1 knockdown on NFkB phosphorylation, IRF3, and IRF7 using phosphoflow. The half overlaid histograms in left panel were generated in Flowjo_v10.8.1 to elucidate the comparison phosphorylated species in control vs si-AL109754.1 transfected cells. After 48h post transfections all the cells were fixed with Cytofix Buffer (BD Biosciences), permeabilized with methanol and stained with various phospho-antibodies for TFs-associated with the TLR signaling. (**B**) Representative scatter plots show the phosphorylated percentage of downstream TLR molecules generated in Flowjo_v10.8.1. (**C**) Bar graph showing average percentage of cells containing phosohorylated TLR specific downstream signaling molecules. Each bar shows mean ±SD. Students t-test were used to calculate p-values and *p*<0.05 was considered significant. *\*p<0.05, **p<0.01, ***p<0.001*.

IRF7 phosphorylation is an important downstream event in the TLR7-mediated activation of Type I IFN pathway (34). Therefore, we have evaluated the phosphorylation of IRF7 in AL109754.1 knockdown DCs post-TLR7 stimulation. Compared to scramble (52.85±2.8%), we observed similar pIRF7 positive cells in AL109754.1 (47.95±10.3%; p<0.05) knockdown DCs. However, the percentage of phosphorylated NF-κB+ cells post- TLR7 stimulation in AL109754.1 knockdown (36.8±5.1%; p<0.05) were remarkably reduced as compared to scramble (46.4±9.4%) (**Fig. 5B, left middle and right panel**). Similar to TLR7, we observed reduced NF-κB phosphorylation when AL109754.1 knockdown DC were stimulated with TLR5 agonist. The percentage of NF-κB phosphorylated cells were remarkably less in AL109754.1 knockdown (49±9.8%; p<0.05) compared to scramble (66±8.4%) (**Fig. 5C, left middle and right panel**). Together these results confirms that reduced expression of TLRs due to knockdown of AL109754.1 also results in reduced TLR-dependent signaling in DCs. Moreover, AL109754.1 confer its regulation of TLR signaling by targeting unique effector molecule upon TLR stimulation.

### LncRNA AL109754.1 knockdown modulates the expression of TLR dependent expression of NF-κB genes

TLR-dependent crosstalk and cross-regulation between IRF3 and NF-κB impacts overall expression of NF-κB regulated genes (33, 34). Importantly, NF-κB orchestrates multitude of genes that eventually regulate innate and adaptive immune activities including the expression of pro-inflammatory cytokines and chemokines genes and regulation of inflammasome functions (35). Thus, in pursuit of the results obtained in previous section, we examined the mechanisms underlying reduced phosphorylation of IRF3, IRF7 and NF-κB by quantifying NF-κB-induced genes in DC transfected with siAL109754.1. Using specific human NF-κB signaling pathway PCR array, we have evaluated the expression of 88 key NF-κB target genes in DCs transfected with control, or siAL109754 upon activation with TLR4, TLR5 or TLR7 agonists.

Compared to control, we observed downregulation of various NF-κB-pathway genes in siAL109754.1 DCs. **Figure 6A-C** show the heatmaps of downregulated genes in AL109754.1 knockdown DCs stimulated with TLR4, 5 and 7 agonists. In TLR4 agonist treated DC, AL109754 knockdown significantly reduced expression of 11 transcripts including EDG2, TICAM1, BIRC4, NF-ΚB1, TNF, NF-ΚB1A, AKT1, ICAM1, BCL2, CD40 and CSF3 (**Figure 6A**). We observed significant downregulation of 15 transcripts viz., IFNA1, NF-ΚB1, ELK1, TLR1, NF-ΚB1A, TNF, SELL, EDG2, EGR1, FOS, HTR2B, CSF2, MMP9, TNRRSF1A and IL12B in siAL109754 knockdown DC stimulated with TLR5 agonists (**Figure 6B**). Furthermore, activation of TLR7 in AL109754 knockdown DC identified significant downregulation of 15 transcripts: CHUK, RHOA, MALT1, TNFRSF10A, FOS, TLR4, BCL10, BIRC4, FADD, JUN, AKT1, CSF1, MMP7, EGR1 and BCL2 (**Figure 6C**). As depicted in Venn diagram, TLR4 and TLR5 stimulation of siAL109754.1 treated DC identified four commonly downregulated genes EDG2, TNF, NF-ΚB1A and ICAM, while TLR4 and TLR7 stimulation showed three common genes BIRC4, BCL2 and AKT1. Comparison of TLR5 and TLR7 data shows common downregulation of EGR1 gene. Intriguingly, none of the genes examined were common to all TLR stimulation, thereby suggesting differential impact of lncRNA AL109754.1 on TLR signaling (**Fig. 6D**). Among the downregulated genes, we noted various cytokines, transcription factors and signaling molecules that integrate TLR signaling, which myeloid corroborates with our previous data and suggesting a broad impact of AL109754.1 potentiating inflammatory phenotype. Our results show that AL109754.1 can regulate unique transcriptional profiles upon TLR activation and innate immune responses by DC.

**FIGURE 6:**
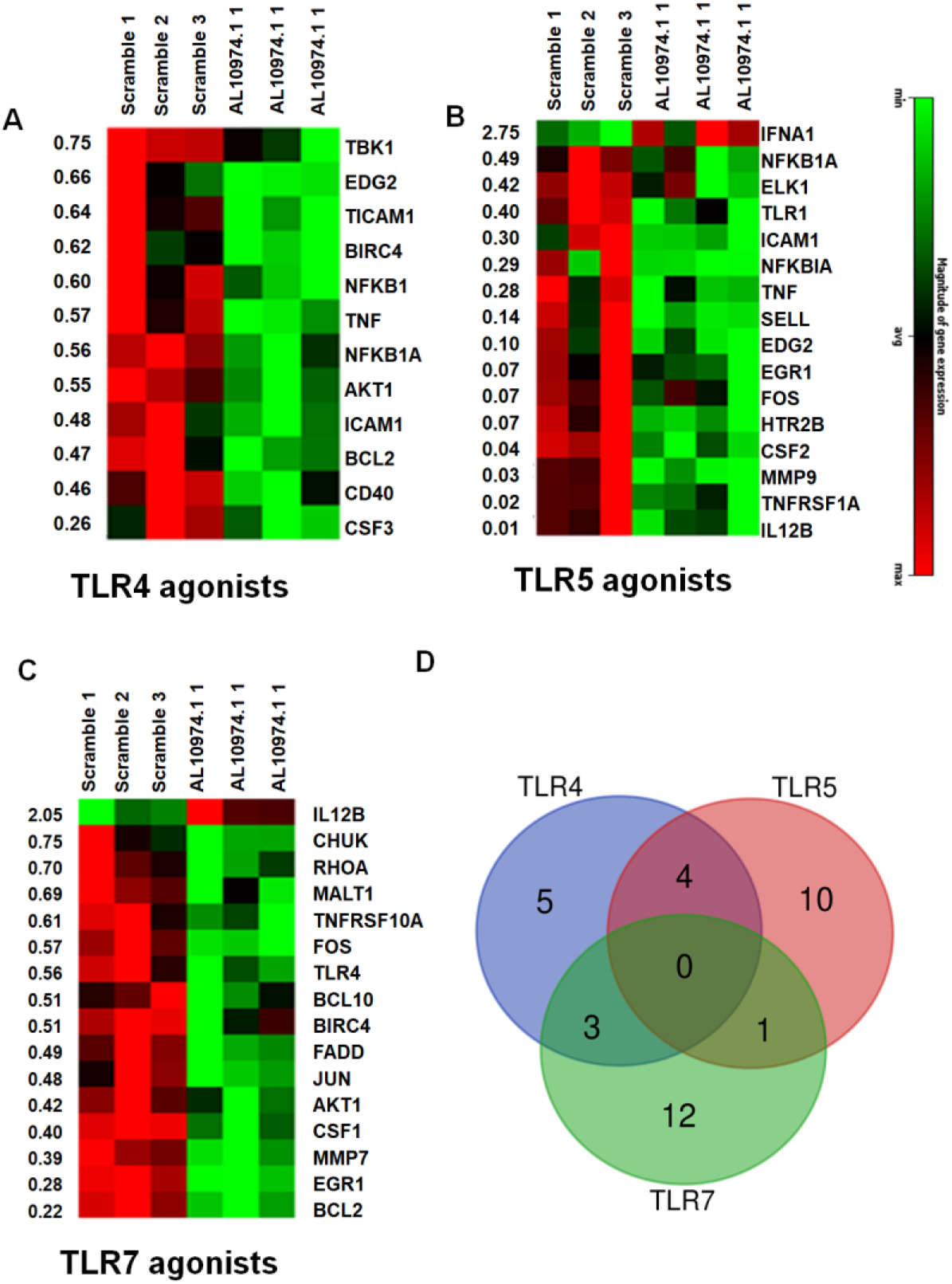
LncRNA AL109754.1 knockdown effects the TLR-directed NFκB dependent gene expression. Scramble or si-AL109754.1 transfected DCs were challenged with TLR4 (concentration), TLR5 (concentration), and TLR7 (concentration) agonists. After 18h post-challenge, total RNA was isolated using miRNeasy Kit (Qiagen). cDNA was synthesized from 500 ng total RNA and the expression of NFκB pathway genes was assessed by custom PCR array containing 88 gene primers. The top of the heat map shows the sample identifiers. Heat maps showing differentially expressed genes in AL109754.1 knockdown DC (n=3/group) challenged with **(A)** TLR4 agonist, **(B)** TLR5 agonist and **(C)** TLR8 agonist. The data was analyzed using the Qiagen GeneGlobe Data Analysis Center (Source: https://www.qiagen.com/us/shop/genesand-pathways/data-analysis-center-overview-page/). B2M was used as housekeeping to normalize the expression levels. The fold change was used to analyze the changes NFκB pathway related genes in DCs challenged with TLR agonists (scramble vs si-AL109754.1; *p<0.05*). (B) Venn diagram showing the overview of differential gene expression of NFκB pathway upon TLR stimulation in AL109754.1 knockdown DCs.

## Discussion

Myeloid dendritic cells (mDCs) is a heterogeneous population of professional antigen-presenting cells that coordinate innate and adaptive immune responses, and immune tolerance (36). Thus, their differentiation and functions should be tightly regulated in order to maintain immune homeostasis, as various high throughput studies have elucidated unique mRNA expression pattern in different DC phenotypes (37). Owing to multitude of regulatory functions orchestrated by lncRNAs viz., transcription factor (TF) recruitment at promoter region and epigenetic reprograming, microRNA sponges, translational regulation of various mRNAs) (38–41), we hypothesized that their unique expression of lncRNA repertoire should drive monocyte-to-DC differentiation process. Thus, we performed a comprehensive time kinetics of lncRNA expressional profiling in monocyte- to DC differentiation using RNA-seq. Our lncRNA datasets obtained in this study is unique and our findings demonstrate contribution of lncRNAs (in terms of up and downregulation) throughout the DC differentiation process. We identified thousands of differentially expressed lncRNAs as early as 18 h post-stimulation.

Interestingly, we report here the stable expression and upregulation of previously unannotated lncRNAs AL109754.1 and AC093278.2 in monocyte-to-DC differentiation, thereby strongly suggesting their key involvement in orchestrating DCs differentiation events. *In vitro* monocyte-to-DC and M1 macrophage differentiation shares GM-CSF as a common agent. Typically, GM-CSF is produced by multiple cells types (macrophages, DCs, T cells, neutrophils, eosinophils, tissue fibroblasts during inflammation (42–43), as proinflammatory cytokines, (IL-1α, IL-1β, TNF-α, and IL-12) are reported to induce GM-CSF, whereas IL-4, IFN-γ, and IL-10 at the site of inflammation (44–48) acts as an essential factor for recruitment, activation and survival of myeloid cells (49–50). The enrichment of AL109754.1 also in M1 rather than M2 macrophages suggest that AL109754.1 might be a signature for expressing in pro-inflammatory microenvironment and required for myeloid inflammatory cell differentiation. Consistent with this, we observed downregulated DC differentiation markers viz., CD1a, CDw93 and CD209 expression in case of AL109754.1 knockdown, but not AC093278.2, strongly suggesting its role in monocyte-to-DC differentiation. This the first report demonstrating the involvement of lncRNA AL109754.1 in monocyte-to-DC differentiation. Previously published reports have identified induction of conserved intergenic lncRNA lnc-DC, and differential expression of 962 lncRNAs in DCs isolated from Allergic rhinitis patients during differentiation of monocyte-to-DCs and in mature DCs, respectively (9, 10). However, all these studies only divulge the fact enlightening the involvement of lncRNAs in DC differentiation, at a particular time, when the data was recorded. In contrast, we observed progressive induction of AL109754.1 expression during DC differentiation while it was absent in the monocytes, precursor cells and the knockdown of AL109754.1 negatively impairs DC differentiation by downregulation of key surface markers..

TLRs expression and signaling is directly linked with DC differentiation and maturation in presence of pathogens (51). Previous studies in this field only embark on TLR activation mediated lncRNA induction or downregulation. Activation of THP1 macrophages with TLR2 (Pam3CSK4) and TLR4 (LPS) agonists results in the induction of Lnc-IL-17R, while other study identified downregulation of lncRNA THRIL in Pam3CSK4-stimulated THP-1 macrophages and induction of cytokine genes upon its ablation in these cells (52). Furthermore, during the course of mice infection studies with Japanese encephalitis virus, rabies (53, 54), IAV, and HSV (55), which induces endosomal TLRs, numerous lncRNAs were dysregulated, including NEAT1. Till now, no clear study suggested the role of lncRNAs in maintaining TLR expression on DC. Contemplating it, we performed knockdown of lncRNA AL109754.1 in DC and extensively evaluated the expression of all TLRs (1–9). Knockdown of AL109754.1 significantly dampens both surface and intracellular TLRs *viz*., TLR2, TLR4, TLR5, TLR7, TLR8 and TLR9, on monocyte-derived DC *in vitro* thereby suggesting that lncRNA expression is needed for the optimal expression of TLRs in DCs. Typically, TLRs located at the cell surface function as PPR for bacteria, whilst intracellular TLR reflect the anti-viral defense. Thus, expression of lncRNA AL109754.1 in DCs likely regulate both anti-bacterial and anti-viral innate immune responses.

Activation of TLRs eventually results in secretion of pro-inflammatory cytokine, which provide adequate microenvironment as well provide signal 3 for the activation of T cells (56). However, in previous studies the role of lncRNAs in regulating cytokine expression is reported in macrophages. Using high-throughput analysis, Li et al. reported the upregulation of Linc1992 in THP1 macrophages and demonstrated it to regulate the secretion of TNFα. linc1992, IL-8, CXCL10, CCL1, and CSF1 (16). This group also showed the upregulation of this lncRNAs in children undergoing treatment and resolution of Kawasaki disease, which was directly correlated with the decreased TNFα levels in the convalescent phase samples (16). Furthermore, reports also demonstrate the induction of lncRNA NeST in patients suffering from immune thrombocytopenia (IT) (57), Sjorgren’s syndrome (58) and ulcerative colitis (59). In this study, we show lncRNA AL109754.2 knockdown significantly attenuates secretion pro-inflammatory cytokines IL-1β, IL-6, IL-8 and TNFα on stimulation of DCs with TLR 2, 4, 5, 7 and 9 agonist strongly suggesting the definitive role of this lncRNA in maintaining TLR expression and their immune activation in DCs. This is the first study that compressively characterized the effect of DC-enriched lncRNA on TLR-expression and the cytokine secretion.

Nuclear translocation and activation of NF-κB, activation of interferon regulatory factors 3/7 (IRF3/7) and/or activator protein-1 (AP-1) are the molecular signatures to recognition of invading pattern recognition receptors (PRRs) by TLRs (Toll-Like Receptors) (25–27). Lately, Zhang et al. demonstrated the genome-wide changes in lncRNA and protein-coding genes in pursuit of TLR signaling in human macrophages. They showed that TLR activation induces multitude of lncRNAs, which are located within the 5 kb of protein coding genes related to immune-related functions, thereby suggesting the TLR mediated immune regulation via expression dynamics of cis acting lncRNA genes (60). Furthermore, Josyln et al. identified 588 TLR7- and IFN-I-dependent lncRNAs and reported their association with multiple processes including dysregulation of pDC maturation, IFN-I and inflammatory cytokine production, antigen presentation, costimulation, tolerance and IFN-I-driven autoimmune diseases such as systemic lupus erythematosus (61). These reports suggest that downstream TLR signaling might regulated by lncRNAs. Studies exploring the lncRNA-mediated control of downstream effector molecules phosphorylation in TLR signaling remain elusive. Herein, we examined the role of AL109754.1 in modulating phosphorylation of IRFs (IRF3 and IRF7) and NF- κB during stimulation of monocyte-to-DC. AL109754.2 knockdown DC challenged with TLR4, TLR7 and TLR5 agonist exhibit significantly reduced phosphorylation of IRF3, IRF7 and NF-κB, further suggesting its role in the activation of TLR signaling. Interestingly, TLR4 stimulation of AL109754.2 knockdown DCs impair IRF3 and IRF7 phosphorylation, not NF-κB phosphorylation. TLR7 agonist disrupts NF-κB phosphorylation not its specific downstream molecule IRF7, whilst TLR5 only effects NF-κB phosphorylation. These results indicate that AL109754.2 regulate crosstalk between TLRs by impairing multiple effector molecule phosphorylation critical in shaping resultant innate immune response in a concerted manner. Whether lncRNAs regulate the IRF phosphorylation is an understudied topic in both antigen presenting cells viz., DC and macrophages. Fan et al., suggested that overexpression of lncATV in huh7 line antagonized hepatitis C-stimulated phosphorylation of TBK1, STAT1, and IRF3 (62). Also, targeted deletion of lncRNA LUCAT1 augments the expression of type I interferon inducible genes in myeloid cells challenged with LPS (63).

To further dissect the role of lncRNA AL109754.2 in TLR activation, we performed focused NF-κB pathway gene PCR array DCs challenged with in TLR4, TLR5 and TLR7 agonist. We observed significant downregulation of multiple genes involved in NF-κB pathway in AL109754.2 knockdown DCs further supporting its role in regulating TLR signaling. Similar to phosphorylation results, we noticed that most of the downregulated genes were unique to each TLR stimulation, few were common to at least two TLRs but none of the genes were common to all three TLRs. These results further demonstrate a key role of AL109754.2 in innate immune activation of myeloid DCs. lncRNA AL109754.2 mediate its functional effects on DC need further studies to decipher the basis of regulation via epigenetic modifications, microRNA control, or protein interaction. Because of the lack of murine homolog of human AL109754.2, further studies should focus on human *in vitro* and *ex vivo* samples to advance the understanding of lncRNA-mediated regulation of immunity in health and disease.

In summary, we profiled global lncRNA profiles of monocyte-differentiated myeloid DCs and identified thousands of differentially expressed lncRNA repertoire. We focused on two lncRNAs that show time-dependent enrichment during the DC differentiation and demonstrate that lncRNA AL109754.2 is an important regulator of DC differentiation, TLR (both surface and intracellular) expression and their immune function (cytokine production) suggesting its role in maintaining immune homeostasis.

## Acknowledgments

We are grateful to NIH/NIDCR R03 DE027147, R01DE027980, and R21DE026259 to AN. We acknowledge Flow cytometry core, UIC for assisting in flow cytometry analyses.

## References

1. Shortman, K., and Y.J. Liu. 2002. Mouse and human dendritic cell subtypes. Nat. Rev. Immunol. 2:151–161.

2. Liu, J., X. Zhang, Y. Cheng, and X. Cao. 2021. Dendritic cell migration in inflammation and immunity. Cell Mol. Immunol. 18: 2461–2471.

3. Ulvmar, M. H., K. Werth, A. Braun, P. Kelay, E. Hub, K. Eller, L. Chan, B. Lucas, I. Novitzky-Basso, K. Nakamura, et al. 2014.The atypical chemokine receptor CCRL1 shapes functional CCL21 gradients in lymph nodes. Nat. Immunol. 15: 623–630.

4. Braun, A. T., Worbs, G. L. Moschovakis, S. Halle, K. Hoffmann, J. Bölter, A. Münk, and R. Förster. 2011. Afferent lymph-derived T cells and DCs use different chemokine receptor CCR7-dependent routes for entry into the lymph node and intranodal migration. Nat. Immunol. 12: 879–887.

5. Ahmad, I., Valverde, A., Ahmad, F., and Naqvi, A.R. 2020. Long Noncoding RNA in Myeloid and Lymphoid Cell Differentiation, Polarization and Function. Cells. 9(2):269.

6. Bocchetti, M., Scrima, M., Melisi, F., Luce, A., Sperlongano, R., Caraglia, M., Zappavigna S., and Cossu, A.M. 2021. LncRNAs and Immunity: Coding the Immune System with Noncoding Oligonucleotides. Int. J Mol. Sci. 22(4):1741.

7. Mattick, J.S., Amaral, P.P., Carninci, P. et al. 2023. Long non-coding RNAs: definitions, functions, challenges and recommendations. Nat. Rev. Mol. Cell Biol. https://doi.org/10.1038/s41580-022-00566-8.

8. Ahmad, I., Valverde, A., Naqvi, R.A., and Naqvi AR. 2020. Long Non-coding RNAs RN7SK and GAS5 Regulate Macrophage Polarization and Innate Immune Responses. Front Immunol. Dec 9;11:604981.

9. Wang, P., Y. Xue, Y. Han, L., Lin, C., Wu, S., Xu, Z., Jiang, J., Xu, Q., Liu, and Cao X. 2014. The STAT3-binding long noncoding RNA lnc-DC controls human dendritic cell differentiation. Science. 344: 310–313.

10. Zhou, Y., Chen, X., Zheng, Y., Shen, R., Sun, S., Yang, F., Min, J., Bao, L., Zhang, Y., Zhao, X., Wang, J., and Wang, Q. 2021. Long Non-coding RNAs and mRNAs Expression Profiles of Monocyte-Derived Dendritic Cells From PBMCs in AR. Front. Cell Dev. Biol. 9:636477.

11. Xin, J., J. Li, Y. Feng, L. Wang, Y. Zhang, and R. Yang. 2017. Downregulation of long noncoding RNA HOTAIRM1 promotes monocyte/dendritic cell differentiation through competitively binding to endogenous miR-3960. Onco Targets Ther. 10:1307–1315.

12. Takeuchi, O., and S. Akira. 2010. Pattern recognition receptors and inflammation Cell. 140: 805–820.

13. Kawasaki, T., and Kawai T. Toll-like receptor signaling pathways. 2014. Front Immunol. 5:461. DOI: 10.3389/fimmu.2014.00461.

14. Zhao, Q., Pang, G., Yang, L., Chen, S., Xu, R., and Shao, W. 2021. Long Noncoding RNAs Regulate the Inflammatory Responses of Macrophages. Cells. Dec 21;11(1):5.

15. Carpenter, S., D. Aiello, M. K., Atianand, E. P., Ricci, P., Gandhi, L. L., Hall, M., Byron, B., Monks, M. Henry-Bezy, J. B. Lawrence et al. 2013. A long noncoding RNA mediates both activation and repression of immune response genes. Science. 341: 789–792.

16. Li, Z., T-C., Chao, K-Y. Chang, N., Lin, V.S., Patil, C., Shimizu, et al. 2014. The long noncoding RNA THRIL regulates TNFα expression through its interaction with hnRNPL. Proc. Natl. Acad. Sci. USA. 111:1002–1007.

17. Böyum A. 1968. Isolation of leucocytes from human blood – further observations. Scand. J. Clin. Lab. Invest. Suppl. 9731–9750.

18. M. Martin. Cutadapt removes adapter sequences from high-throughput sequencing reads. EMBnet j. 17:10-. DOI: 10.14806/ej.17.1.200.

19. Langmead B., C. Trapnell, M. Pop, and S. L. 2009. Salzberg Ultrafast and memory-efficient alignment of short DNA sequences to the human genome. Genome Biol.10:R25.

20. 20. Kim, D., Pertea, G., Trapnell, C. et al. 2013. TopHat2: accurate alignment of transcriptomes in the presence of insertions, deletions and gene fusions. Genome Biol 14, R36. https://doi.org/10.1186/gb-2013-14-4-r36.

21. Pertea, M., D. Kim, G.M. Pertea, J. T. Leek, and S. L. Salzberg. 2016. Transcript-level expression analysis of RNA-seq experiments with HISAT, StringTie and Ballgown. Nat. Protoc. 11:1650–1667.

22. Kong L, Zhang Y, Ye ZQ, Liu XQ, Zhao SQ, Wei L, and Gao, G. 2007. CPC: assess the protein-coding potential of transcripts using sequence features and support vector machine. Nucl. Acid. Res. 35:W345–9.

23. Atianand, M.K., and Fitzgerald K. A. 2014. Long non-coding RNAs and control of gene expression in the immune system. Trends Mol. Med. 20:623–631.

24. Fordham, J. B., Naqvi, A. R., and Nares, S. 2015. Regulation of miR-24, miR-30b, and miR-142-3p during macrophage and dendritic cell differentiation potentiates innate immunity. J Leukoc. Biol. 98:195–207.

25. de Saint-Vis, B., Vincent, J., Vandenabeele, S., Vanbervliet, B., Pin, J.J., Aït-Yahia, S., Patel, S., Mattei, M.G., Banchereau, J., Zurawski, S., Davoust, J., Caux, C., and Lebecque, S. 1998. A novel lysosome-associated membrane glycoprotein, DC-LAMP, induced upon DC maturation, is transiently expressed in MHC class II compartment. Immunity. 9:325–336.

26. Akira, S., K. Takeda, and T. Kaisho. 2001. Toll-like receptors: critical proteins linking innate and acquired immunity. Nat. Immunol. 2: 675–680.

27. Jarrossay, D., Napolitani, G., Colonna, M., Sallusto, F. and Lanzavecchia, A. 2001. Specialization and complementarity in microbial molecule recognition by human myeloid and plasmacytoid dendritic cells. Eur. J Immunol. 11: 3388–3393.

28. Fang, R., Wang, C., Jiang, Q., Lv, M., Gao, P., Yu, X., Mu, P., Zhang, R., Bi, S., Feng, J. M. and Jiang, Z. 2017. NEMO-IKKβ Are Essential for IRF3 and NF-κB Activation in the cGAS-STING Pathway. J Immunol. 199:3222–3233.

29. Hemmi H., and Akira, S. 2005. TLR signalling and the function of dendritic cells. Chem. Immunol. Allergy. 86:120–135.

30. Jarrossay, D., Napolitani, G., Colonna, M., Sallusto, F., and Lanzavecchia, A. 2001. Specialization and complementarity in microbial molecule recognition by human myeloid and plasmacytoid dendritic cells. Eur. J Immunol. 11: 3388–3393.

31. Kawai, T., and Akira. S. 2010. The role of pattern-recognition receptors in innate immunity: update on Toll-like receptors. Nat. Immunol. 11:373–384.

32. McGettrick, A. F., and O’Neill, L. A. 2010. Localization and trafficking of Toll-like receptors: an important mode of regulation. Curr. Opin. immunol. 22:20–27.

33. Fitzgerald, K. A., Rowe, D. C., Barnes, B. J, Caffrey, D. R., Visintin, A, Latz, E., Monks, B., Pitha, P. M., and Golenbock, D. T. 2003. LPS-TLR4 signaling to IRF-3/7 and NF- kappaB involves the toll adapters TRAM and TRIF. J Exp. Med. 198:1043–1055.

34. Doyle, S., Vaidya, S., O’Connell, R., Dadgostar, H., Dempsey, P., Wu, T., Rao, G., Sun, R., Haberland, M., Modlin, R,. and Cheng, G. 2002. IRF3 mediates a TLR3/TLR4-specific antiviral gene program. Immunity. 17:251–263.

35. Xing, Y., Cao, R., and Hu, H.M. 2016. TLR and NLRP3 inflammasome-dependent innate immune responses to tumor-derived autophagosomes (DRibbles). Cell Death Dis. 7:e2322.

36. Suss, G. and Shortman, K. A 1996. subclass of dendritic cells kills CD4 T cells via Fas/Fas-ligand induced apoptosis. J. Exp. Med. 183, 1789–1796.

37. Alcántara-Hernández, M., Leylek, R., Wagar, L.E., Engleman, E.G., Keler, T., Marinkovich, M.P., Davis, M. M., Nolan, G.P., and Idoyaga, J. 2017. High-Dimensional Phenotypic Mapping of Human Dendritic Cells Reveals Interindividual Variation and Tissue Specialization. Immunity. 47(6):1037–1050.e6.

38. Rossi, M.N., and Antonangeli, F. LncRNAs: New Players in Apoptosis Control. Int. J. Cell Boil. 2014, 2014, 1–7.

39. Marchese, F.P., Raimondi, I., and Huarte, M. 2017. The multidimensional mechanisms of long noncoding RNA function. Genome Biol. 2017, *18*, 206.

40. Dhanoa, J.K., Sethi, R.S., Verma, R., Arora, J.S., and Mukhopadhyay, C.S. 2018. Long non-coding RNA: Its evolutionary relics and biological implications in mammals: A review. J. Anim. Sci. Technol. 60, 25. 60:25. doi: 10.1186/s40781-018-0183-7.

41. Salviano-Silva, A. Lobo-Alves, S.C., De Almeida, R.C., Malheiros, D., and Petzl-Erler, M.L. 2018. Besides Pathology: Long Non-Coding RNA in Cell and Tissue Homeostasis. Non-Coding RNA, 4, 3. ; doi: 10.3390/ncrna4010003.

42. Bagby G. C. Jr., Dinarello C. A., Wallace P., Wagner C., Hefeneider S., and McCall E. 1986. Interleukin 1 stimulates granulocyte macrophage colony-stimulating activity release by vascular endothelial cells. J. Clin. Invest. 78:1316–1323.

43. Gasson J. C. 1991. Molecular physiology of granulocyte-macrophage colony-stimulating factor. Blood 77: 1131–1145.

44. Lukens J. R., Barr M. J., Chaplin D. D., Chi H., and Kanneganti T. D. 2012. Inflammasome-derived IL-1β regulates the production of GM-CSF by CD4(+) T cells and γδ T cells. J. Immunol. 188: 3107–3115.

45. El-Behi M., Ciric B., Dai H., Yan Y., Cullimore M., Safavi F., Zhang G. X., Dittel B. N., Rostami A. 2011. The encephalitogenicity of T(H)17 cells is dependent on IL-1- and IL- 23-induced production of the cytokine GM-CSF. Nat. Immunol. 12: 568–575.

46. Jansen J. H., Wientjens G. J., Fibbe W. E., Willemze R., Kluin-Nelemans H. C. 1989. Inhibition of human macrophage colony formation by interleukin 4. J. Exp. Med. 170: 577–582.

47. Ozawa H., Aiba S., Nakagawa, and Tagami H. 1996. Interferon-gamma and interleukin-10 inhibit antigen presentation by Langerhans cells for T helper type 1 cells by suppressing their CD80 (B7-1) expression. Eur. J. Immunol. 26: 648–652.

48. Sagawa K., Mochizuki M., Sugita S., Nagai K., Sudo T., and Itoh K. 1996. Suppression by IL-10 and IL-4 of cytokine production induced by two-way autologous mixed lymphocyte reaction. Cytokine 8: 501–506.

49. Ponomarev E. D., Shriver L. P., Maresz K., Pedras-Vasconcelos J., Verthelyi D., and Dittel B. N. 2007. GM-CSF production by autoreactive T cells is required for the activation of microglial cells and the onset of experimental autoimmune encephalomyelitis. J. Immunol. 178: 39–48.

50. Parajuli B., Sonobe Y., Kawanokuchi J., Doi Y., Noda M., Takeuchi H., Mizuno T., Suzumura A. 2012. GM-CSF increases LPS-induced production of proinflammatory mediators via upregulation of TLR4 and CD14 in murine microglia. J. Neuroinflammation 9: 268.

51. Duan, T., Du, Y., Xing, C., Wang, H.Y., and Wang, R. F. 2022. Toll-Like Receptor Signaling and Its Role in Cell-Mediated Immunity. Front Immunol.13:812774. doi: 10.3389/fimmu.2022.812774.

52. Murphy, M. B., and Medvedev, A. E. 2016. Long noncoding RNAs as regulators of Toll-like receptor signaling and innate immunity. J Leukoc Biol. 99:839–850.

53. Saha, S., Murthy, S., and Rangarajan, P. N. 2006. Identification and characterization of a virus-inducible non-coding RNA in mouse brain. J. Gen. Virol. 87: 1991–1995.

54. Murphy, M. B, and Medvedev, A. E. 2016. Long noncoding RNAs as regulators of Toll-like receptor signaling and innate immunity. J Leukoc. Biol. 99:839–50.

55. Imamura, K., Imamachi, N., Akizuki, G., Kumakura, M., Kawaguchi, A., Nagata, K., Kato, A., Kawaguchi, Y., Sato, H., Yoneda, M., Kai, C., Yada, T., Suzuki, Y., Yamada, T., Ozawa, T., Kaneki, K., Inoue, T., Kobayashi, M., Kodama, T., Wada, Y., Sekimizu, K., Akimitsu, N. 2014. Long noncoding RNA NEAT1-dependent SFPQ relocation from promoter region to paraspeckle mediates IL8 expression upon immune stimuli. Mol. Cell 53, 393–406.

56. Cen, X., and Liu, S., Cheng K. 2018. The Role of Toll-Like Receptor in Inflammation and Tumor Immunity. Front. Pharmacol. 9: 878. doi: 10.3389/fphar.2018.00878.

57. Li, H., Hao, Y., Zhang, D., Fu, R., Liu, W., Zhang, X., Xue, F., Yang, R. 2016. Aberrant expression of long noncoding RNA TMEVPG1 in patients with primary immune thrombocytopenia. Autoimmunity. 49: 496–502.

58. Padua, D. M., Mahurkar-Joshi, S., Law, I. K. M., Polytarchou, C., Vu, J. P., Pisegna, J. R., Shih, D. Q., Iliopoulos, D., Pothoulakis, C. 2016. A long noncoding RNA signature for ulcerative colitis identifies IFNG-AS1 as an enhancer of inflammation. Am. J Physiol. Gastrointest. Liver Physiol. 311: G446–G457.

59. Wang, J., Peng, H., Tian, J., Ma, J., Tang, X., Rui, K., Tian, X., Wang, Y., Chen, J., Lu, L., et al. 2016. Upregulation of long noncoding RNA TMEVPG1 enhances T helper type 1 cell response in patients with Sjögren syndrome. Immunol. Res. 64: 489–496.

60. Zhang, Q., Chao, T. C., Patil, V. S., Qin, Y., Tiwari, S. K., Chiou, J., Dobin, A., Tsai, C. M., Li, Z., Dang, J., Gupta, S., Urdahl, K., Nizet, V., Gingeras, T. R., Gaulton, K. J., Rana TM. 2019. The long noncoding RNA ROCKI regulates inflammatory gene expression. EMBO J. 38 :e100041. doi: 10.15252/embj.2018100041.

61. Joslyn, R. C., Forero, A., Green, R., Parker, S. E., Savan, R. 2018. Long Noncoding RNA Signatures Induced by Toll-Like Receptor 7 and Type I Interferon Signaling in Activated Human Plasmacytoid Dendritic Cells. J Interferon Cytokine Res. 38:388–405.

62. Fan, J., Cheng, M., Chi, X., Liu, X., Yang, W. 2019. A human long non-coding RNA LncATV promotes virus replication through restricting RIG-I-Mediated innate immunity. Front. Immunol. 10. doi: 10.3389/fimmu.2019.01711.

63. Agarwal, S., Vierbuchen, T., Ghosh, S., Chan, J., Jiang, Z., Kandasamy, R. K., Ricci, E., Fitzgerald, K. A. 2020. The long non-coding RNA LUCAT1 is a negative feedback regulator of interferon responses in humans. Nat Commun. 11:6348. doi: 10.1038/s41467-020-20165-5.

