## Supplemental Table 1 and 2 for "Long noncoding RNA AL109754.1 Regulates Myeloid Dendritic Cell Differentiation and Potentiates TLR signaling"

**Supplemental Tables**

Supplemental Table I: Primers used to validate lncRNAs in monocyte and DCs.

| lncRNA | Forward Primer (5’ to 3’) | Reverse Primer (5’ to 3’) |
| --- | --- | --- |
| AL109754.1 | AGC TGC TTC CTG TCC ATT GT | TCA TGC CTC CTG GGT ATT TC |
| AC093278.2 | CCT GGG ATG ATC GTG GTT TA | GCC TTC TGG TTC ATC AGC TC |

Supplemental Table II: List of DsiRNA to silence lncRNAs.

| lncRNA | DsiRNA sequence |
| --- | --- |
| AL109754.1 | 5’-rCrUrUrCrArUrGrArGrCrCrArArCrUrGrArA-3’  5’-rArCrUrGrUrUrUrUrUrCrArGrUrUrGrGrCrU-3’ |
| AC093278.2 | 5’-rCrUrUrUrUrUrArCrArUrUrArUrArCrUrArG-3’  5’-rUrCrArUrCrUrUrCrUrArGrUrArUrArArUrG-3’ |
