## Supplemental Figure 1, 2, 3 for "Long noncoding RNA AL109754.1 Regulates Myeloid Dendritic Cell Differentiation and Potentiates TLR signaling"

### Supplementary Figures

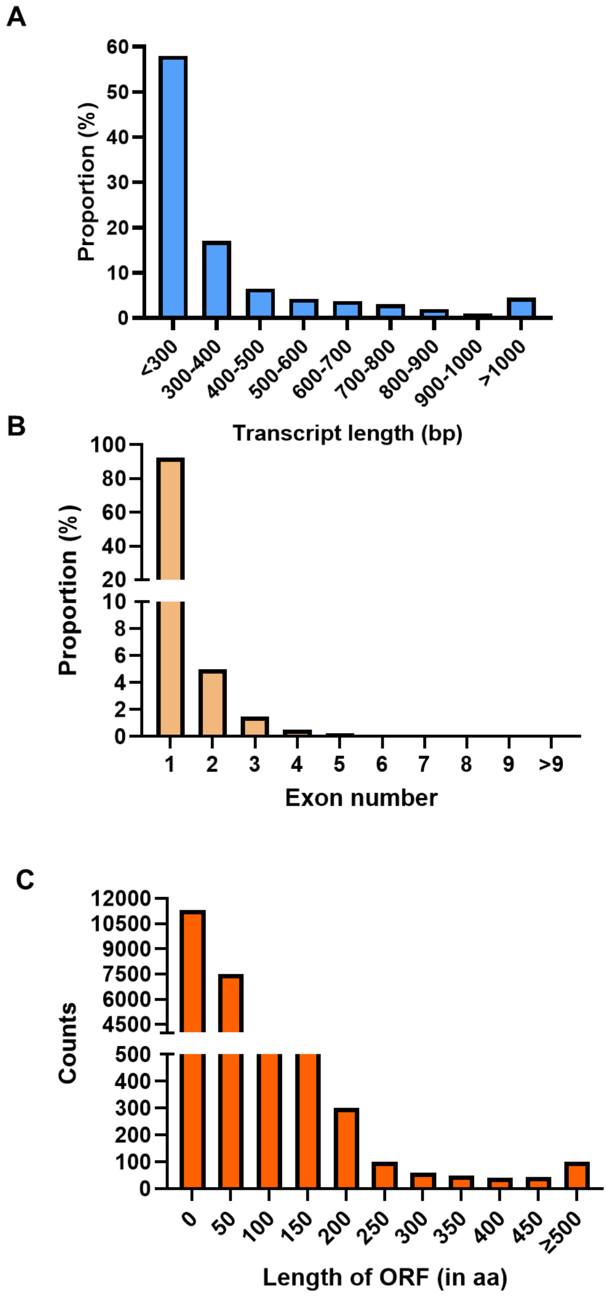


**Supplemental Figure S1.** Histogram showing overall **(A)** percentage of lncRNA transcript length, **(B)** percentage of exon numbers for each lncRNA and **(C)** length of ORF in amino acids (aa) identified in our datasets.

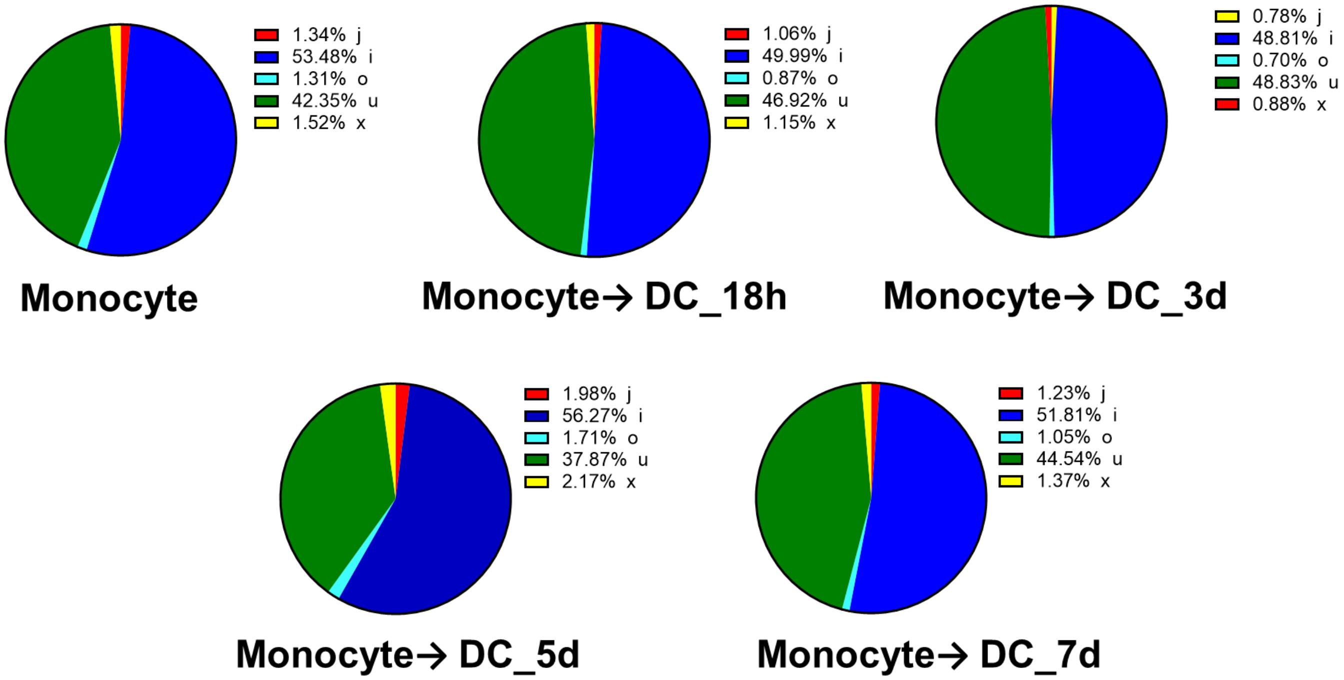


**Supplemental Figure S2.** Various classes of lncRNAs identified in RNA-Seq data during monocytes-to-DC transition. Based on the lncRNAs class code generated by StringTie, lncRNAs were categorized in following different classes: (i) a transfrag located entirely within a reference intron (intronic); (j) potentially novel isoform or fragment with at least one splice junction shared with reference transcript; (o) generic exonic overlap with a reference transcript; (u) unknown, intergenic transcript (intergenic); (x). Pie charts showing percentage of each lncRNA class in each sample.

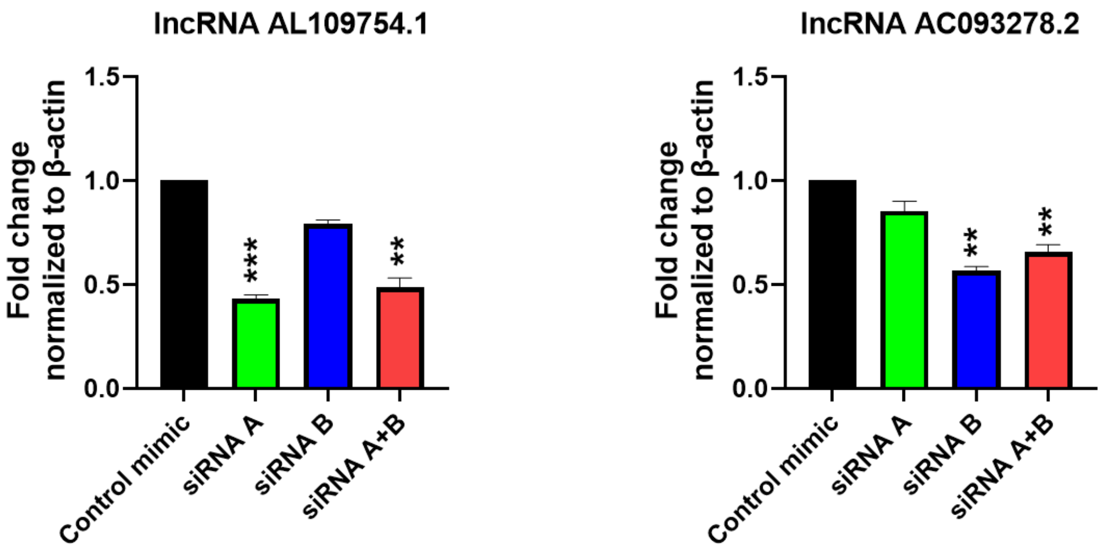


**Supplemental Figure S3**. Testing of siRNAs to knockdown the expression of candidate lncRNAs. DCs were transfected with control, individual siRNA or their combination and the expression of target transcript was quantified by RT-qPCR after 48 h. Each bar shows mean ±SD. Students t-test were used to calculate P-values and P <0.05 was considered significant. *p<0.05, **p<0.01, ***p<0.001.
